# Evaluation of folliculin detection by immunohistochemistry in Birt-Hogg-Dubé associated kidney tumors

**DOI:** 10.1101/2022.06.01.494402

**Authors:** Iris E. Glykofridis, Irma van de Beek, Wim Vos, Pim C. Kortman, Paul van de Valk, Raimundo Freire, Arjan C. Houweling, Rob M.F. Wolthuis

## Abstract

Germline inactivating mutations in folliculin (FLCN) cause Birt–Hogg–Dubé (BHD) syndrome, a rare autosomal dominant disorder predisposing to kidney tumors. Kidney tumors associated with BHD typically lack FLCN expression due to loss of heterozygosity. In this study we assessed the potential of four commercial anti-FLCN antibodies for immunohistochemistry, as currently no routine diagnostic FLCN stainings are performed in the clinic. Despite comprehensive testing, we could not identify a commercial anti-FLCN antibody that is reproducibly effective in immunohistochemical analyses of formalin-fixed paraffin-embedded tissue material. We propose that dedicated future efforts are required to develop a suitable antibody for diagnostic immunohistochemical stainings. The inclusion of FLCN expression status as part of standard renal tumor pathology may contribute to better analyses of the molecular pathology of BHD tumors and facilitate identification of BHD patients, improve their (genetic and clinical) counseling, and enable genetic testing of at risk relatives.

## Introduction

In Birt-Hogg-Dubé (BHD) syndrome, mono-allelic germline pathogenic variants in the folliculin (*FLCN*) gene predispose to an increased risk of bilateral and multifocal renal tumorigenesis [1, 2]. In BHD patients, loss of heterozygosity, by gene silencing or an inactivating somatic mutation of the wild-type *FLCN* allele, is crucial for the development of renal tumors [3-5]. Although different histological renal cell carcinoma (RCC) subtypes have been reported in BHD, there is an overrepresentation of the (hybrid oncocytic-)chromophobe subtype [6-8].

Usually, the diagnosis of RCC can be established by comprehensive histologic examination and a limited immunohistochemistry panel, consisting of markers to distinguish the specific subtypes [9]. Immunohistochemical characterization of 32 BHD-associated renal tumors revealed that in both BHD-associated and sporadic RCCs the immunostaining patterns were comparable[10]. The (hybrid oncocytic-)chromophobe subtypes showed positivity for cytokeratin 7 (*CK7*), kidney-specific cadherin, E-cadherin, CD82 and S100-A1 and negativity for carbonic anhydrase IX (*CA-IX*). BHD-associated and sporadic clear-cell RCCs show CA-IX positivity and CK7 negativity, whereas papillary RCCs stain positive for α-methylacyl-CoA racemase. Despite their histological resemblance at first sight, BHD-associated chromophobe RCCs exhibit distinct molecular characteristics and lack mutations in driver genes such as *TP53* and *PTEN*, which are present in 32% and 9% of sporadic chromophobe RCCs respectively[11]. Whole-exome sequencing analysis of 29 histologically diverse BHD-associated RCCs revealed that copy number variations in these tumors are considerably different from those already reported in sporadic cases [12]. Very few common somatic variants were observed in BHD-associated RCCs, though variants in several chromatin remodeling genes were observed 59% of BHD-associated RCCs.

FLCN protein expression status in renal tumors is not routinely analyzed by immunohistochemistry for diagnostic purposes, although absence of FLCN protein expression may be a strong indication for BHD syndrome. Previous studies performed FLCN immunohistochemical stainings of one BHD-associated renal tumor using a custom-made antibody [13, 14]. Although loss of FLCN protein expression is expected in most BHD-tumor tissues, the immunohistochemical analysis showed robust staining [13, 14]. Apart from the germline mutation, the renal tumor of this particular BHD patient had a *FLCN* splice-site variant c.397-7_404del15. This variant was predicted to result in exon skipping and production of a truncated FLCN protein, which could be an explanation for the positive staining in the tumor tissue.

Here, we comprehensively tested four commercially available anti-FLCN antibodies to assess their applicability for immunohistochemical analyses of FLCN protein expression in formalin-fixed paraffin-embedded (FFPE) renal tissue material. In addition, we performed tests with our new custom-made anti-FLCN antibody. The inclusion of sensitive and specific FLCN protein expression status in standard renal tumor pathology is likely to contribute to increased identification of BHD patients and improve their (genetic) counseling.

## Materials & methods

### Cell culture and CRISPR mediated gene editing

Renal proximal tubular epithelial cells (RPTEC/TERT1, ATCC® CRL-4031™) were maintained in DMEM/F12 (Gibco ®, Life Technologies) according to the manufacturer’s protocol with addition of 2% fetal bovine serum (FBS, Gibco ®, Life Technologies). To maintain the selective pressure for immortalization 0.1mg/mL G418 Sulfate (Calbiochem) was added. CRISPR/Cas9 mediated gene editing was used to disrupt FLCN expression as previously described [15]. In short, RNP complexes (Cas9 protein + Synthego FLCN_exon 4 GAGAGCCACGAUGGCAUUCA + modified EZ scaffold) were transfected transiently in RPTEC cells using Neon Electroporation System (ThermoFisher), where after knock-out status of single-cell derived clones was validated by western blot and Sanger sequencing.

BHD renal tumor cell line UOK257 and its FLCN-reconstituted version UOK257-2 [16, 17] were maintained in DMEM (Gibco ®, Life Technologies) with 8% FBS (FBS, Gibco ®, Life Technologies). To maintain the selective pressure for FLCN expression in UOK257-2, medium was supplemented with 2 μg/ml Blasticidin (Invitrogen, Life Technologies).

All cell lines were cultured in a humidified atmosphere at 37°C and 5% CO_2_ and were regularly tested to exclude Mycoplasma infections.

### Immunoblotting

Cell line lysates were made in NP40 lysis buffer (50 mM Tris-HCl pH 7.4, 150 mM NaCl, 1% NP40) supplemented with protease inhibitors (Roche). Next, samples were boiled at 70°C for 5 min in 1x NuPAGE LDS sample buffer (Novex NP0007, Thermo-Fischer) with 10% 1M DTT (Sigma) and equal amounts were separated by 4–15% or 8–16% SDS-PAGE (BioRad) and blotted onto polyvinylidene fluoride (PVDF) transfer membranes (Merck). Subsequently, membranes were blocked for 1 hr at room temperature with 5% milk (ELK, Campina, Amersfoort, Netherlands) or Bovine Serum Albumin (BSA) in TBST. The primary anti-FLCN antibody incubations were overnight at 4°C in 2.5% milk in TBST. The next day, membranes were washed and incubated with appropriate secondary antibodies (Dako) for 3 hr at 4°C in 2.5% milk or BSA in TBST. After incubation, the membranes were thoroughly washed where after bands were visualized by chemiluminescence (ECL prime, Amersham, VWR, Radnor, Pennsylvania, USA) in combination with ChemiDoc Imaging Systems (BioRad, Hercules, California, USA).

Anti-FLCN antibodies were used in following dilutions: Ab #1 (1:1000 in milk, #3697, Cell Signaling Technology), Ab #2 (1:500 in milk, ab93196, Abcam), Ab #3 (1:1000 in BSA, HPA028760, Atlas antibodies) and Ab #4 (1:1000 in BSA, ab176707, Abcam). Anti-β-Actin (c-47778, Santa Cruz) and anti-vinculin (1:1000, sc-25336, Santa Cruz) antibodies were used according to individual datasheets.

### Immunocytochemistry

For immunocytochemistry cell pellets (∼15E6 cells) were fixed in 4% paraformaldehyde for a minimum of 4 hours. Fixed cells were resuspended in liquid agar and placed at 4°C to solidify. Next, cell-agar pieces were embedded in paraffin to obtain a cytoblock, that was subsequently used for sectioning and antibody stainings.

### Immunohistochemistry

For all tissue material used in this study, there was consent for research purposes. Paraffin sections of 4-μm thickness were subjected to immunohistochemistry. After deparaffinization and rehydration, antigen retrieval was mediated by heat (autoclaved at 121°C for 20 minutes) or pepsin treatment (at 37°C for 10 minutes). Then, samples were treated with the diluted anti-FLCN antibodies for 1 hour. Different combinations of buffers and antigen retrieval methods and dilutions were tested, of which the most promising ones are depicted below. Immunohistochemical stainings were done using the EnVision-Detection System/HRP (Dako, Glostrup, Denmark), followed by 3,3-diaminobenzidine (DAB+) chromogen staining. Heamatoxylin was used for counterstaining of nuclei, where after slides where washed with tap water and mounted with cover slips.

| **Antibody** | **Material** | **Dilution** | **Buffer** | **Antigen retrieval** |
| --- | --- | --- | --- | --- |
| <u>Ab #1</u><br>CST #3692, D14G9 lot 3 | cell lines | 1:100 | 10 mM Tris/1mM EDTA-buffer, pH 8.5 | heat-mediated (HIER) |
| <u>Ab #2</u><br>Abcam ab93196 | cell lines | 1:100 | 0,01M citrate buffer pH6 | heat-mediated (HIER) |
| <u>Ab #3</u><br>Atlas antibodies<br>HPA028760 | cell lines | 1:100 | 10 mM Tris/1mM EDTA-buffer, pH 8.5 | heat-mediated (HIER) |
| <u>Ab #4</u><br>Abcam ab176707 | cell lines | 1:100 | Tris buffer pH 2.0 | Pepsin-mediated<br>(PIER) |
| <u>Ab #1</u><br>CST #3692, D14G9 lot 3 | tissues | 1:25 | 10 mM Tris/1mM EDTA-buffer, pH 8.5 | heat-mediated<br>(HIER) |
| <u>Ab #3</u><br>Atlas antibodies<br>HPA028760 | tissues | 1:25 | 0,01M citrate buffer pH6 | heat-mediated<br>(HIER) |
| <u>Custom-made Ab</u> | cell lines | 1:1000 | 10 mM Tris/1mM EDTA-buffer, pH 9 | heat-mediated<br>(HIER) |

The custom-made antibody was raised in rabbits immunized with recombinant protein containing amino acids 280-580 of the human FLCN. To obtain the antigen, the corresponding cDNA fragment of human *FLCN* was cloned in the pET-30 vector (Novagen) and the His-tagged fusion protein was expressed in *E. coli* and was purified using Ni-NTA (Qiagen) before injection. For affinity purification of the sera, around 150 μg of the *FLCN* antigen was loaded onto SDS–PAGE and transferred to a nitrocellulose membrane. After staining with Ponceau S (Merck-Millipore), the part of the membrane with the antigen was cut out, blocked with 2% BSA in TBS-T for 1 h and then incubated with 3 ml serum overnight at 4 °C. Bound antibodies were eluted with 0.15 M glycine-HCl, pH 2.3 and after elution Tris-HCl, pH 8.8, was added to reach pH of the antibody solution to pH 7.5. For validations of the custom-made antibody, full-length *FLCN* was cloned into pEGFP-C1 and pEGFP-N1 expression plasmids, which were subsequently transfected in U2OS cells (supplementary figure 1).

## Results & discussion

An overview of the four, commercially available anti-FLCN antibodies tested in this study is shown in table 1, along with their specific characteristics such as resource, host, clonality and specific antigen.

**Table 1.** Overview and characteristics of commercial anti-FLCN antibodies tested

| Antibody | Resource | Host + clonality | Antigen |
| --- | --- | --- | --- |
| #1 | Cell Signaling Technology (#3697) | Rabbit mAb | Total protein |
| #2 | Abcam (ab93196) | Rabbit pAb | Center region<br>sequence proprietary |
| #3 | Atlas antibodies (HPA028760) | Rabbit pAb | C-terminal<br>VGCEDDQSLSKYEFV<br>VTSGSPVAADRVGPTI<br>LNKIEAALTQNLSVD<br>VVD<br>QCLVCLKEEWMNKVK<br>VLFKFTKVDSRPKEDT<br>QKLLSILGASEEDNVK<br>LLK<br>FWMTGLSKTYKSHLM<br>STVR |
| #4 | Abcam (ab176707) | Rabbit pAb | N-terminal<br>MNAIVALCHFCELHG<br>PRTLFCTEVLHAPLPQ<br>GDGNEDSPGQGEQA<br>EEEEG |

To investigate the specificity of these four antibodies, we started with western blot analyses of FLCN wild type (FLCN^POS^) and FLCN deficient (FLCN^NEG^) cell line lysates. As shown in figure 1A, the four different antibodies were first tested on lysates of renal proximal tubular epithelial cells (RPTEC/TERT1) in which CRISPR/Cas9 mediated genome editing was used to endogenously knock-out FLCN [15]. Except antibody #2, all antibodies revealed a specific FLCN band at the predicted height, which was lacking in the FLCN^NEG^ condition. Antibody #1 was the only antibody showing no additional back ground bands. Repeating the same experiment with another isogenic cell line pair, the BHD renal tumor derived cell line (UOK257, FLCN^NEG^) and its FLCN-reconstituted version (UOK257-2, FLCN^POS^) revealed a similar pattern (Figure 1B). As we eventually want to assess the potential of these antibodies in immunohistochemistry, we proceeded with testing the specificity of the four antibodies in immunocytochemistry. For this, FLCN^POS^ and FLCN^NEG^ UOK257 cell line pellets were fixed and embedded in paraffin for subsequent sectioning and staining. The results of the immunocytochemistry stainings are shown in figure 1C. Only antibodies #1 and #3 showed an expected result, with clear staining in the FLCN^POS^ condition and absence of staining in the FLCN^NEG^ condition. However, in FLCN^NEG^ cells stained with antibody #1 there appeared to be some nuclear signal present. For antibodies #2 and #4 there was a clear signal in both conditions, which makes these antibodies not suitable for further immunohistochemistry purposes.

**Figure 1.**
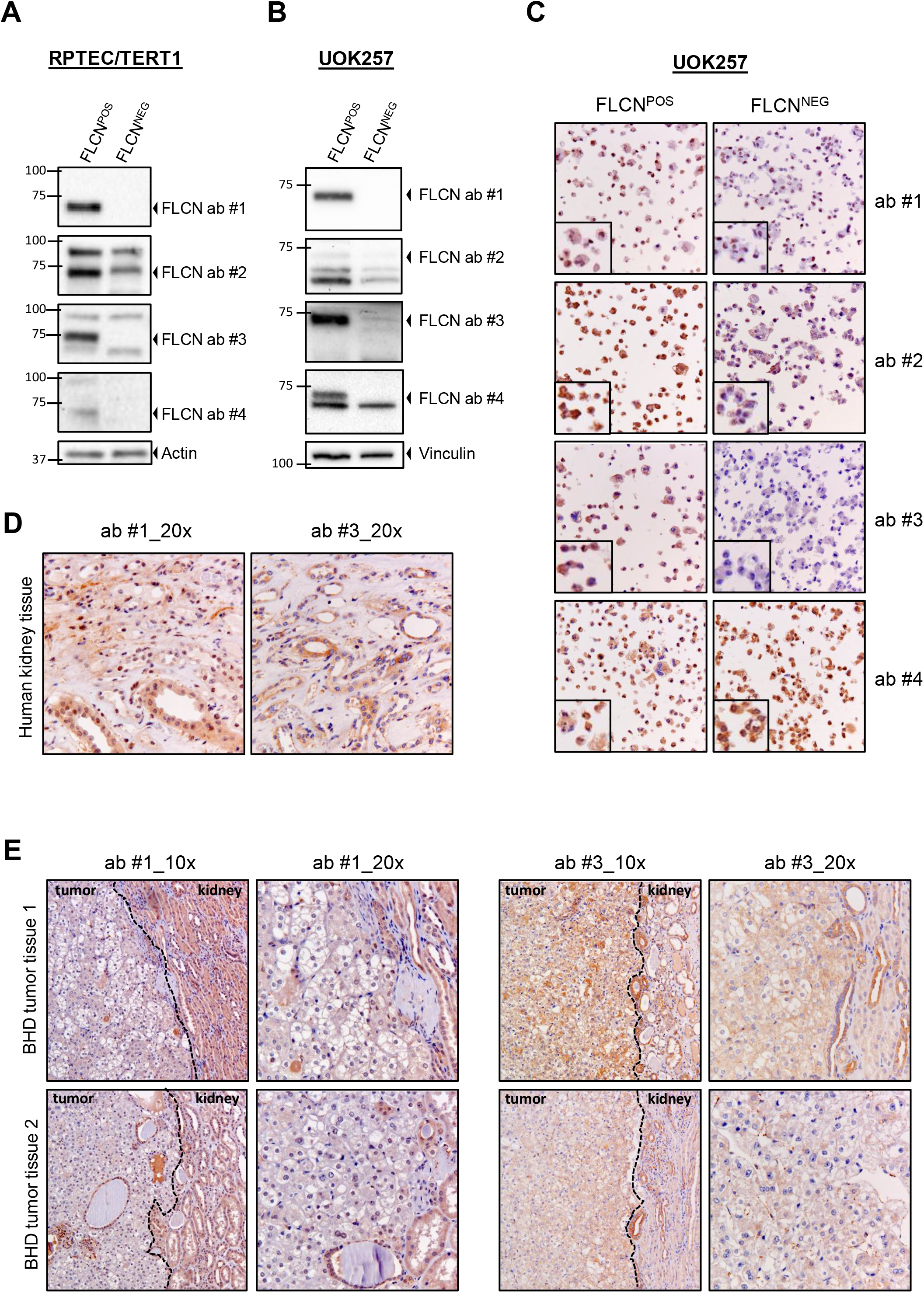
Evaluation of the specificity and efficacy of commercial antibodies to detect FLCN by western blot and immunohistochemistry. **A**. Western blot of wild type and FLCN knock-out RPTECs using four different, commercial anti-FLCN antibodies. Actin was used as a loading control. **B**. Western blot of BHD kidney tumor cell line UOK257 (FLCN^NEG^) and the same cell line with FLCN expression restored UOK257-2 (FLCN^POS^). Four different, commercial anti-FLCN antibodies were used. Vinculin was used as a loading control. **C**. Immunocytochemistry of UOK257 cell lines (FLCN^POS^ vs. FLCN^NEG^) using four different, commercial anti-FLCN antibodies. Images were taken with 10x objective and zoom-ins are shown in lower left corner of each staining. Brightness and contrast adjustments were applied to the whole image. **D**. Immunohistochemistry of human kidney tissue using anti-FLCN antibodies #1 and #3. Images were taken with 20x objective. Brightness and contrast adjustments were applied to the whole image. **E**. Immunohistochemistry of two BHD renal tumor tissues, using anti-FLCN antibodies #1 and #3. Images were taken with 10x and 20x objectives. The border of tumor and surrounding tissue is indicated with a line on images with lower magnification. Brightness and contrast adjustments were applied to the whole image.

As a next step, we proceeded with antibodies #1 and #3 and performed FLCN immunohistochemistry analyses of normal human kidney tissue obtained from the Amsterdam UMC BioBank (Figure 1D). Both antibodies showed a clear staining pattern, with strong signal in the tubular cells specifically. Then we assessed whether these antibodies could be used to demonstrate loss of FLCN expression in tumor tissues, as is the case in most Birt-Hogg-Dubé syndrome associated renal tumors. We obtained two BHD-associated chromophobe RCC FFPE samples from the Amsterdam UMC BioBank and performed immunohistochemical stainings with anti-FLCN antibodies #1 and #3 (Figure 1E). The tumor tissues were derived from BHD patients with a c.499C>T (tissue 1) and c.319_320delGTinsCAC (tissue 2) germline variant in *FLCN*, respectively. The borders of tumor and surrounding tissue are indicated with lines on photos with a lower magnification, indicating that especially with antibody #1 there is absence of staining in the tumor area, while in the surrounding tissue the staining is present. This indicates that FLCN protein expression is indeed lost in the tumor. For antibody #3 the difference in signal between tumor and surrounding tissue was less distinct, but at higher magnifications it was clear that for many cells in the tumor area staining was absent.

Taken together, we concluded that anti-FLCN antibody #1 had the best potential to be used for immunohistochemistry purposes and proceeded with the collection of additional BHD-associated renal tumors to evaluate its efficacy in a larger sample batch. However, when a more recent lot number of the antibody was used, the staining signals appeared to be less strong and specific, while all other protocol conditions were kept identical. For this reason, we decided that the chance to use this antibody for routine FLCN stainings for pathological purposes was low and we did not proceed to use it in further experiments.

In parallel, we performed additional analyses of the antibody directed against the C-terminus, raised against amino acids 522–536 of human FLCN, that was described previously [13, 14]. Before, no *FLCN* knock-out control experiments were included and antibody specificity validations were not shown. Unfortunately we were not able to obtain a FLCN-specific signal when testing this antibody in our cell line models. The BHD-tumor tissue previously used for FLCN immunohistochemistry, was the same as our BHD-tumor tissue 1 (Figure 1E). Remarkably, in contrast to the clear positive staining previously observed, we detect no positive signal in the tumor area when using anti-FLCN antibody #1. This difference could be caused by the fact that antibody #1 was raised against total F protein. However, our antibody #3 was raised against a C-terminal peptide as well (amino acids 452-570) and indeed did show a positive signal in the particular tumor too, albeit weak. Therefore, we cannot exclude that this BHD tumor retains expression of a truncated FLCN protein as a result of the specific splice-site variant.

Additionally, we generated an anti-FLCN antibody ourselves, raised against amino acids 280-580 of human FLCN, and performed multiple experiments to assess its specificity and performance in immunocytochemistry. A western blot of FLCN^POS^ and FLCN^NEG^ UOK257 cell line lysates (supplementary figure 1A) showed that the affinity-purified custom-made antibody recognizes the FLCN protein specifically. As shown in supplementary figure 1B, we could only observe a strong, specific signal in cells transfected with a FLCN-expression construct. This indicates that the antibody is specific, but that the low endogenous expression levels of FLCN further hamper the detection of this protein in tissues.

A recent study by Zhao and colleagues [18] showed that the FLCN protein is marginally expressed in human clear cell renal cell carcinoma tissues using a novel commercial anti-FLCN antibody (from ProteinTech) for their immunohistochemistry experiments. This antibody may be included in future studies to validate whether it is also suitable for diagnostic purposes, i.e. the antibody is specific and has a constant quality over different batches. Meanwhile, we are carrying out additional attempts to produce a high affinity anti-FLCN antibodies for immunohistochemistry. Notably, given the results shown in Figure 1, it may also be considered to develop an anti-FLCN western or dot-blotting strategy of BHD tumor material for primary clinical diagnosis, followed by genetic testing.

The inclusion of FLCN expression status as part of standard renal tumor pathology may contribute to increased identification of BHD patients and improve their (genetic) counseling. Alternatively, indirect biomarkers for loss of FLCN expression in renal cells may be used for (early) diagnosis of BHD patients. Also, with the recent improvements in speed and price of sequencing technologies, it could be an option to perform *FLCN* genotype testing of every RCC patient identified, which is currently gaining momentum for germline variants that predispose to breast cancer [19].

---

**Supplementary figure 1.**
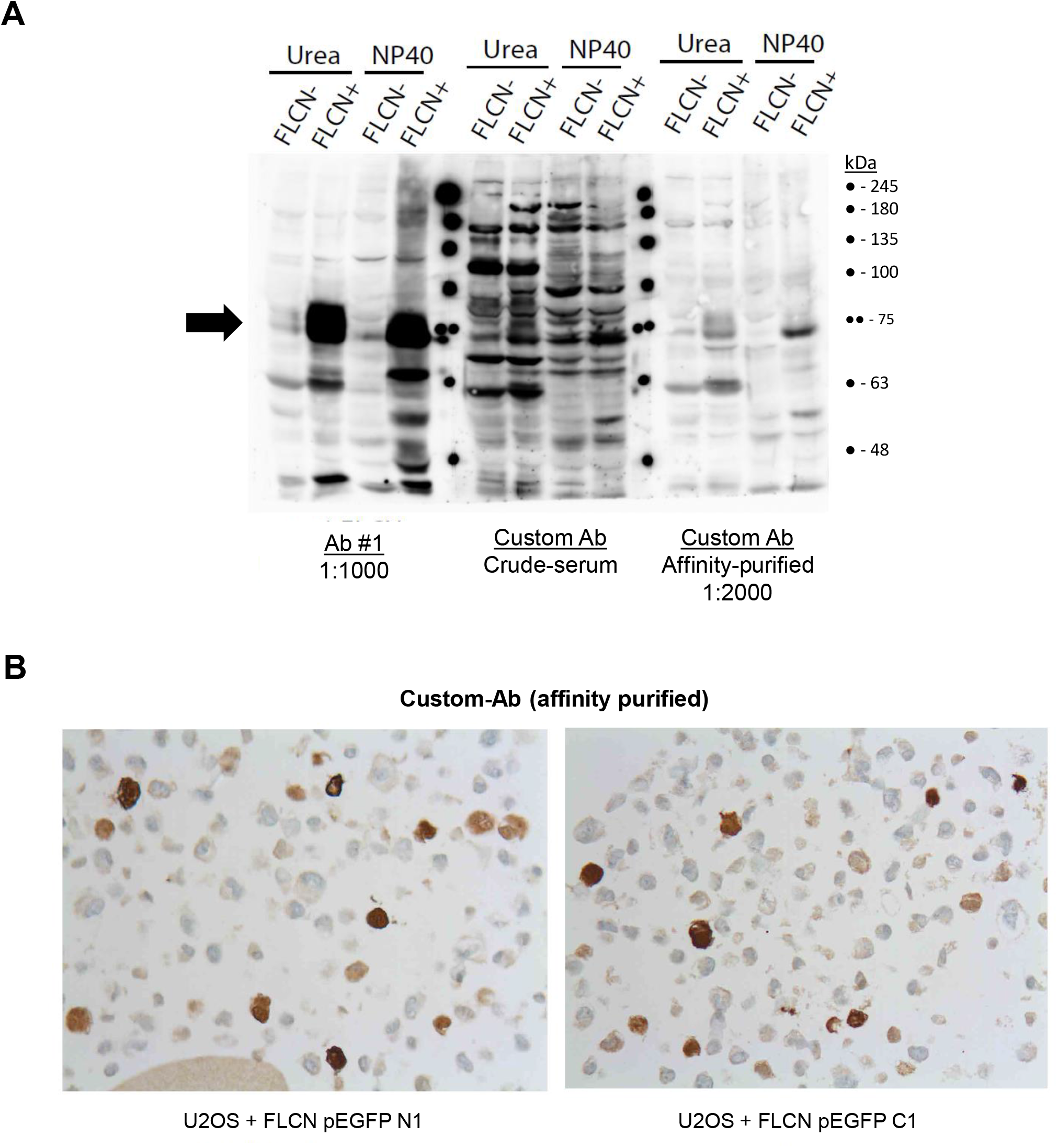
Evaluation of the specificity and efficacy of a custom made antibody to detect FLCN by western blot and immunocytochemistry. **A**. Western blot of UOK257 (FLCN-) and UOK257-2 (FLCN+) cell line lysates shows that Ab #1 and the affinity-purified custom-made Ab do recognize the FLCN protein specifically. The arrow indicates the predicted height, based on the size of the FLCN protein (∼64kDa). **B**. Immunocytochemistry of U2OS cell lines transfected with FLCN-GFP expression constructs (N1 =N-terminally tagged and C1=C-terminally tagged). Only cells that have high expression show a strong positive signal. Images were taken with 20x objective.

